# Compensatory functional connectome changes in a rat model of traumatic brain injury

**DOI:** 10.1101/2021.05.17.444382

**Authors:** Zhihui Yang, Tian Zhu, Marjory Pompilus, Yueqiang Fu, Jiepei Zhu, Kefren Arjona, Rawad Daniel Arja, Matteo M. Grudny, H. Daniel Plant, Prodip Bose, Kevin K. Wang, Marcelo Febo

## Abstract

Penetrating cortical impact injuries alter neuronal communication beyond the injury epicenter, across regions involved in affective, sensorimotor, and cognitive processing. Understanding how traumatic brain injury (TBI) reorganizes local and brain wide nodal functional interactions may provide valuable quantitative parameters for monitoring pathological progression and functional recovery. To this end, we investigated spontaneous fluctuations in the functional magnetic resonance imaging (fMRI) signal obtained at 11.1 Tesla in rats sustaining controlled cortical impact (CCI) and imaged at 2- and 30-days post-injury. Graph theory-based calculations were applied to weighted undirected matrices constructed from 12,879 pairwise correlations between fMRI signals from 162 regions. Our data indicate that on days 2 and 30 post-CCI there is a significant increase in connectivity strength in nodes located in contralesional cortical, thalamic, and basal forebrain areas. Rats imaged on day 2 post-injury had significantly greater network modularity than controls, with influential nodes (with high eigenvector centrality) contained within the contralesional module and participating less in cross-modular interactions. By day 30, modularity and cross-modular interactions recover, although a cluster of nodes with low strength and low eigenvector centrality remain in the ipsilateral cortex. Our results suggest that changes in node strength, modularity, eigenvector centrality, and participation coefficient track early and late TBI effects on brain functional connectivity. We propose that the observed compensatory functional connectivity reorganization in response to CCI may be unfavorable to brain wide communication in the early post-injury period.

## Introduction

Traumatic brain injury (TBI) is a leading cause of emergency department visits, long-term disability, and accidental deaths in the U.S. (CDC, 2019). Among different types of moderate-to-severe brain injury, penetrative concussive cortical injury can cause permanent structural and functional deficits in brain areas that regulate cognitive, sensorimotor, and affective functions (Arciniegas, 2011; Hegde, 2014; Johnstone *et al*., 2015; Thomas *et al*., 2015). The severity of cognitive impairment and/or neuropsychiatric sequelae of focal cortical injury may be linked to the degree of spatial damage caused at and beyond the TBI epicenter (Hall *et al*., 2005). Degeneration of axons and dendrites can occur beyond the TBI foci and produce impairments in brain wide communication across cortical and subcortical regions (Hall *et al*., 2005; Hall *et al*., 2008; Carron *et al*., 2016; Ping and Jin, 2016).

The large-scale neurobiological changes that occur with TBI have been investigated using resting state functional magnetic resonance imaging (fMRI) and graph theory-based analysis of functional connectivity (Hillary *et al*., 2011; Sharp *et al*., 2011). Disruption of functional connectivity between subdivisions of the caudate and putamen and distributed cortical regions, including the anterior cingulate, was observed to be associated with cognitive impairments in a cohort of 42 TBI patients compared to aged-matched control participants (De Simoni *et al*., 2018). Structural connectivity-based network matrices in 52 TBI patients, 21 of which sustained microbleeds, were shown to have a reduction in nodes with high betweenness and eigenvector centrality in caudate and anterior cingulate cortex (Fagerholm *et al*., 2015). These nodes are highly influential within networks, constituting routes of high traffic or communication between distant regions of the cortex. Cognitive processing was closely linked to the degree of disconnection in network hubs, perhaps because of a reduced number of influential nodes following diffuse axonal injury (Fagerholm *et al*., 2015). A comprehensive longitudinal blast related TBI study reported significant modular organizational changes (Han *et al*., 2014). Functional connectivity can be parcellated into groups or modules with greater rates of within-group connectivity than between-group node interactions. This modular pattern is thought to be a fundamental organizational aspect of neural activity in the brain of various species and may be important in the segregation of functions across distributed networks. The number of between-module connections, as quantified by the participation coefficient, is reduced with blast injury and this has important implications for cognitive function and the binding of multiple sources of sensory, affective, and memory information.

The above neuroimaging studies in human subjects emphasize the promise of graph theory measures as functional TBI biomarkers. Comparable methods in animal models of TBI allow investigation of neurobiological mechanisms linked to network metric changes. Repetitive closed head injury disrupts functional connectivity between midbrain, hippocampal, and cerebellar regions and produced hyperconnectivity in olfactory regions and other areas of the rat brain (Kulkarni *et al*., 2019), and caused widespread changes in tissue apparent diffusion coefficient values (Yang *et al*., 2015). Niskanen and colleagues provided evidence of post-concussive functional recovery of BOLD signal responses to forepaw electrical stimulation in primary somatosensory (S1) cortex by day 56 in a lateral fluid percussive injury (LFPI) rat model (Niskanen *et al*., 2013). Rats sustaining LFPI were also shown to have reduced cortical functional connectivity at four months post-injury (Mishra *et al*., 2014). In a series of studies using the controlled cortical impact (CCI) model in rats, Harris and colleagues demonstrated contralesional increases in cortical excitability, c-Fos expression, and BOLD signal responses to forepaw electrical stimulation (Verley *et al*., 2018). Interestingly, the S1 cortical hyperexcitability was observed as early as 2 days post-injury using electrophysiological methods but at later time points with fMRI data collected at 7 Tesla (Verley *et al*., 2018). Consistent with these findings, further functional connectomic assessments by Harris and colleagues demonstrated increased network strength and topological reorganization involving interactions between strongly and weakly connected regions (Harris *et al*., 2016).

In the present study, we investigated functional connectivity changes in a rat CCI model at 2- and 30-days post-injury. Our results indicate that at two days post-injury cortical regions contralateral to the CCI epicenter show a significant increase in node strength along with a global increase in modularity, with low and high centrality nodes largely distributed in different modules. Participation in cross-modular connectivity was also reduced on day 2, however, this recovered to control levels by day 30 post-CCI.

## Materials and Methods

### Subjects

Female and male Sprague-Dawley rats (220-300g) were obtained from Charles River Laboratories (Raleigh, NC, USA.). Rats were housed in sex-matched pairs in a temperature and humidity-controlled vivarium with a 12 h light cycle (lights on at 0700 h) and food and water provided ad libitum. Rats were assigned to one of two experimental conditions: controls (15 male and 8 female rats) and controlled cortical impact (CCI; 19 male and 12 female rats). A subgroup of CCI rats were imaged 2 days post-injury (16 male and 7 female rats) and another group (3 male and 5 female rats) at 30 days. All procedures received prior approval from the Institutional Animal Care and Use Committee of the University of Florida and followed all applicable NIH guidelines.

### Controlled cortical impact (CCI)

The CCI procedure was carried out using a Leica Impact One device (Leica Microsystems Inc). Anesthetic levels were induced with 4% isoflurane gas mixed in 100% oxygen and the maintained under 2% isoflurane for the rest of the procedure. Body temperature was regulated to 37°C using a thermal pad while rats were prepared for surgery on a stereotaxic frame. A parasagittal craniectomy (center at anteroposterior [AP], +4.0 mm; L, +2.8 mm from lambda) 5 mm in diameter was performed to expose the brain and allow impactor tip access to the cortical surface. The impactor had a 4-mm flat-face tip. CCI at a depth of 1.5 mm at 4 m/s and a dwell time of 240ms was carried out. All injuries occurred in the right hemisphere. The surgical area was sutured, and recovery was monitored by tail pinch and righting reflexes.

### Magnetic resonance imaging

Magnetic resonance imaging was carried out as previously reported (Pompilus *et al*., 2020). Images were collected in an 11.1 Tesla MRI scanner (Magnex Scientific Ltd., Oxford, UK) with a Resonance Research Inc. gradient set (RRI BFG-240/120-S6, maximum gradient strength of 1000 mT/m at 325 Amps and a 200 µs risetime; RRI, Billerica, MA) and controlled by a Bruker Paravision 6.01 console (Bruker BioSpin, Billerica, MA). A custom-made 2.5 cm x 3.5 cm quadrature radiofrequency (RF) surface transmit/receive coil tuned to 470.7MHz (^1^H resonance) was used for B1 excitation and signal detection (RF engineering lab, Advanced Magnetic Resonance Imaging and Spectroscopy Facility, Gainesville, FL).

Rats were scanned under a continuous flow of 1.5 % isoflurane (delivered at 0.5L/min mixed with medical grade air containing 70% N2 and 30% O2). Respiratory rates were monitored continuously, and body temperature was maintained at 36-37°C using a warm water recirculation system (SA Instruments, Inc., New York). For each rat, we acquired a 10-minute high-resolution T2 weighted anatomical scan followed by a 10-minute functional magnetic resonance imaging (fMRI) scan. A T2-weighted Turbo Rapid Acquisition with Refocused Echoes (TurboRARE) sequence was acquired with the following parameters: effective echo time (TE) = 41 ms, repetition time (TR) = 4 seconds, RARE factor = 16, number of averages = 12, field of view (FOV) of 24 mm x 18 mm and 0.9 mm thick slice, and a data matrix of 256 × 192 and 25 interleaved ascending coronal (axial) slices covering the entire brain from the rostral-most extent of the anterior frontal cortical surface, caudally towards the upper brainstem and cerebellum. Functional images were collected using a single-shot spin-echo echo planar imaging (EPI) sequence with the following parameters: TE = 15 ms, TR = 2 seconds, 300 repetitions, FOV = 24 × 18 mm and 0.9 mm thick slice, and a data matrix of 64 × 48 with 25 interleaved ascending coronal slices in the same space as the T2 anatomical. Respiratory rates, isoflurane concentration, body temperature, lighting, and room conditions were kept constant across subjects.

### Image pre-processing and atlas registration

The image pre-processing steps applied in the present study are illustrated in the schematic in Figure 1. Anatomical and functional scan masks outlining rat brain boundaries were generated in MATLAB using Three-Dimensional Pulsed Coupled Neural Networks (PCNN3D) (Chou *et al*., 2011). Resting state processing was carried out using software tools in Analysis of Functional NeuroImages (AFNI) (Cox, 1996), FSL (Jenkinson *et al*., 2002), and Advanced Normalization Tools (ANTs) (Klein *et al*., 2009). First, we used 3dDespike in AFNI to remove time series spikes and 3dvolreg for image volume alignment. Preprocessed scans were cropped and a high-pass temporal filter (>0.009Hz) was used (3dTproject) to remove slow variations (temporal drift) in the fMRI signal. Independent component analysis (ICA) decomposition was then applied to preprocessed scans to assess noise components in each subjects’ native space prior to spatial smoothing and registration. The resulting components were assessed, and in most cases all components contained noise-related signal along brain edges, in ventricular voxels, and large vessel regions. These components were suppressed using a soft (‘non-aggressive’) regression approach, as implemented in FMRIB Software Library (FSL 6.0.3) using fsl_regfilt (Jenkinson *et al*., 2002). A low-pass filter (<0.12Hz) and spatial smoothing (0.6mm FWHM) was then applied to the fMRI scans.

**Figure 1.**
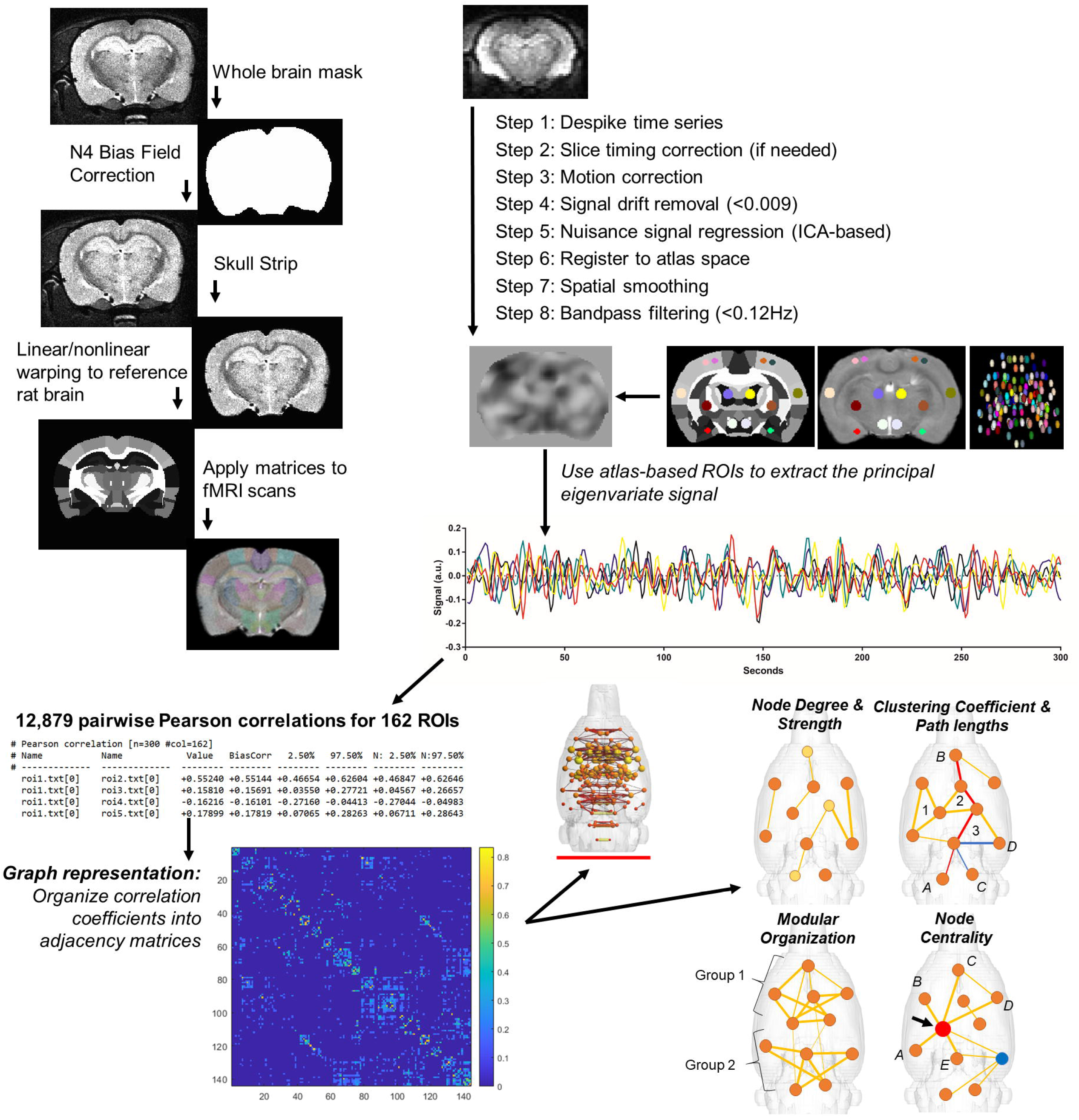
Image processing and analysis workflow used in the present study. Upper left (top to bottom) shows anatomical scan to atlas registration steps. Upper right shows resting state fMRI pre-processing steps and signal extraction based on 1mm diameter spherical seeds. Bottom panels (left to right) show symmetric matrices of pairwise correlations of fMRI signals from 81 bilateral regions of interest (162 ROI total). Networks are visualized using atlas-based ROI coordinates in a 3-dimensional translucent rat brain shell with vertex sizes represented by spheres and weighed by node strength, and edges as lines weighed by correlation coefficients. Schematic of measures of network and nodal integration, segregation, and centrality that were derived from constructed weighted matrices.

Preprocessed anatomical and fMRI scans were aligned to a parcellated rat brain template (Kenkel *et al*., 2016). Anatomical scans were cropped and N4 bias field correction (Tustison *et al*., 2010) was applied to remove B1 RF field inhomogeneities and reduce field of view intensity variations (Kenkel *et al*., 2016). The extracted brain maps were then linearly registered to the rat template using FSL linear registration tool (FLIRT) (Jenkinson *et al*., 2002), using a correlation ratio search cost, full 180-degree search terms, 12 degrees of freedom and trilinear interpolation. The linear registration output was then nonlinearly warped to the template space using ANTs (antsIntroduction.sh script). The resulting deformation field images were used to generate Jacobian determinant maps to evaluate subject-to-atlas registration quality and quantify the effects of regional structural differences of CCI rats compared to controls. Average log-normalized Jacobian maps are shown in Figure 2. Anatomical-to-atlas linear and nonlinear transformation matrices were applied to fMRI scans at a later stage.

**Figure 2.**
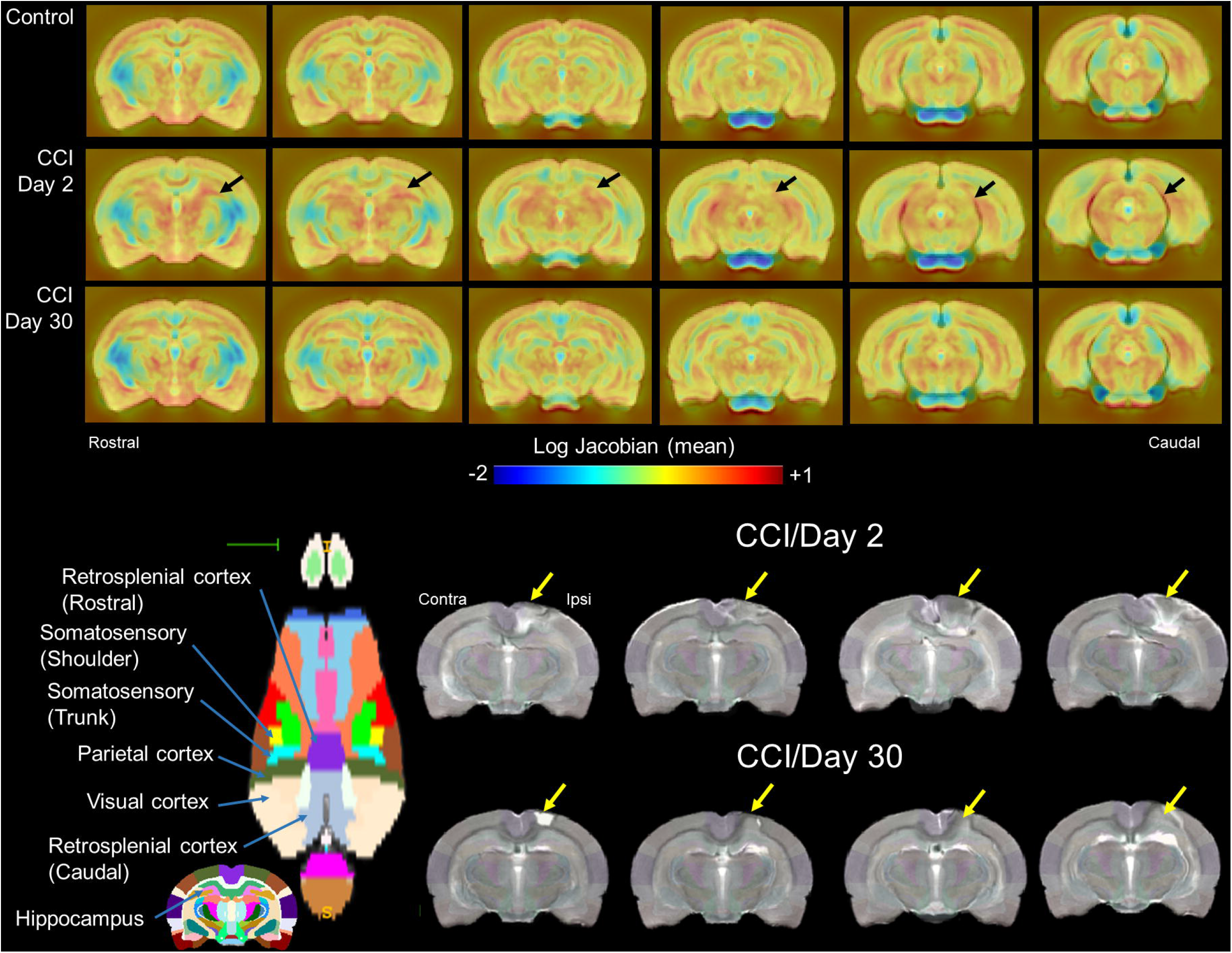
Nonlinear registration to a reference T2-weighted scan revealed cortical and subcortical thalamic structural differences in controlled cortical impact (CCI) rats relative to controls. Top row shows log-normalized Jacobian maps scaled between −2 to +1 (blue to red; narrowing and expansive differences, respectively). Jacobian maps represent the average of each experimental group for controls (n=23), CCI day 2 (n=23) day 30 (n=8) post-injury. Bottom rows show axial view of cortical parcellation highlighting brain areas impacted by CCI. Also shown are four representative anatomical scans of CCI rats acquired at 2- and 30-days post-injury.

Brain extraction using a mask (see above) was first applied to fMRI scans and the cropped scans were then aligned to their respective higher resolution anatomical scans. Timeseries functional images were split into 300 individual volumes and the first in the series was linearly aligned to the anatomical scan using FLIRT (same parameters as above, except 6 degrees of freedom was used in this step). ANTs (antsRegistrationSyNQuick.sh script) was used to warp the lower resolution functional images to their structural (using a single stage step deformable b-spline syn with a 26-step b-spline distance). Linear and nonlinear warping matrices for fMRI-to-anatomical alignment were applied to individual scans in the time series, then the merged 4-D functional timeseries were moved to the atlas space using the prior anatomical-to-template transformation matrices.

### Parcellation-based node placement and resting state signal extraction

We updated a previous set of 144 node parcellations (Pompilus *et al*., 2020) to include an additional 18 region of interest (ROI) masks covering voxels in the CCI region and the contralateral hemisphere (Figure 3A). These included 4 ROIs covering the primary visual cortex (Lesion ROI1, 3-5 in Figure 3A), one in the dorsal hippocampus centered on the dentate gyrus layer (Lesion ROI2), the rostral and caudal retrosplenial cortex (Lesion ROI9 and ROI6, respectively), and the barrel field and trunk regions of the primary somatosensory cortex (Lesion ROI7 and ROI8, respectively). Thus, a total of 162 region of interest (ROI) masks, divided into 81 left and 81 right ROI’s, were included in our analyses. Individual ROI masks were generated using a previously published rat brain parcellation (Kenkel *et al*., 2016). ROI seed generation was as previously reported (Pompilus *et al*., 2020) and coordinates were used for 3D network visualizations in BrainNet viewer in MATLAB (Xia *et al*., 2013). The principal eigenvector timeseries vector was extracted from preprocessed fMRI scans with the assistance of ROI mask overlays. This generated 162 individual ROI text files per subject that contained L2-normalized resting state signals as a vector of 300 data points. The timeseries files were used in cross-correlations and in calculations of Pearson r coefficients for every pairwise combinations of ROIs (1dCorrelate in AFNI). The resulting number of pairwise correlations was 12,879 per subject (after removing 162 self-correlations). Correlation coefficients were imported to MATLAB and Fisher’s transform applied to ensure a normal distribution of z values prior to analyses.

**Figure 3.**
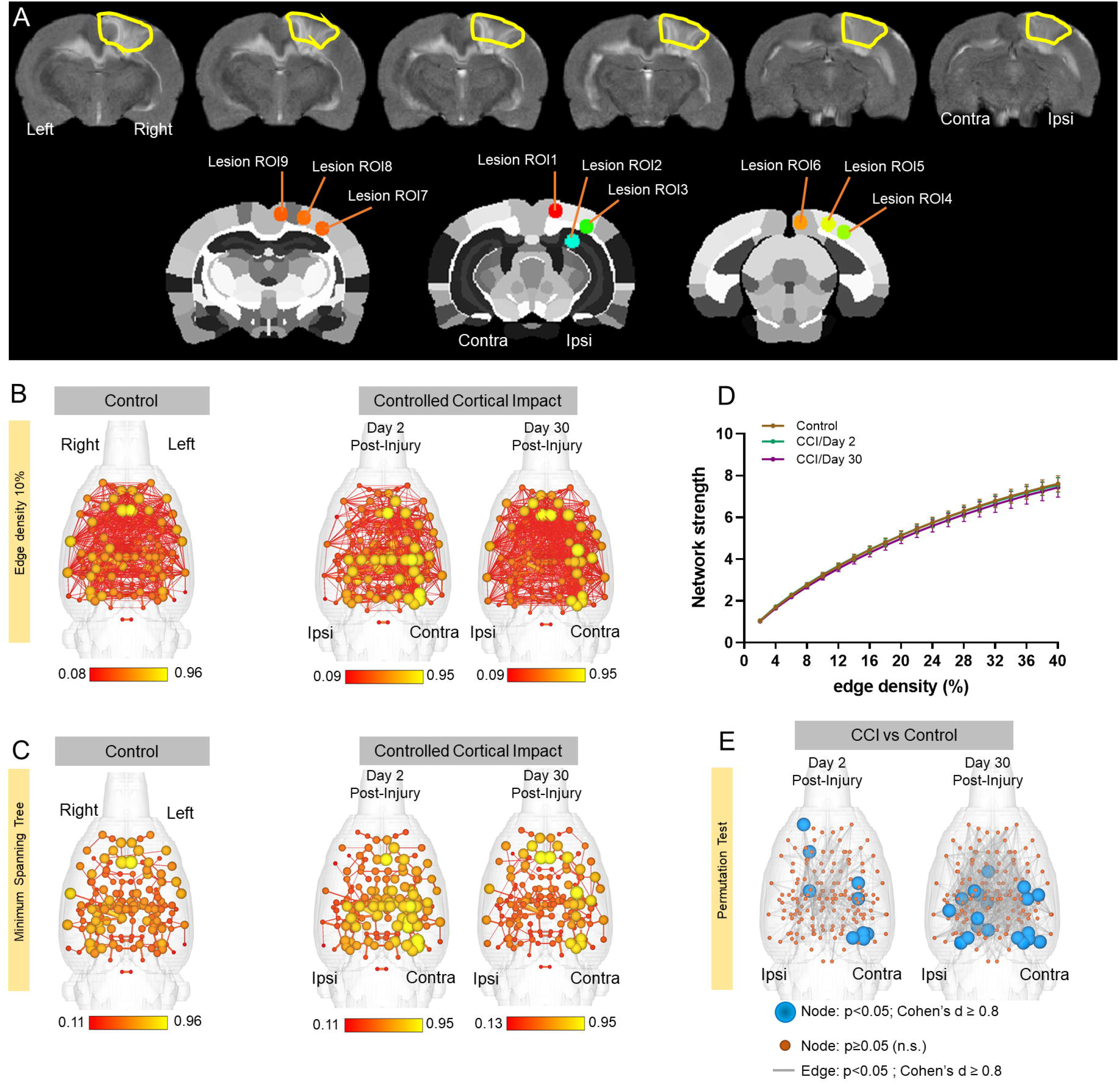
Nodes in cortical regions contralateral to the controlled cortical impact (CCI) site show a significant increase in connectivity strength at day 2 and 30 post-injury. A) A representative T2 anatomical scans highlighting the extent of cortical injury with CCI in the right hemisphere. Lesion ROIs were generated in CCI regions to capture resting state signals from the injury site and homologous contralateral representations. B) 3D functional connectome maps at 10% edge density for control and CCI rats imaged 2- and 30-days post-injury. Scale bars are edge weights. C) Minimum spanning tree for same networks in B, illustrating strongest weighted pairwise correlations. D) Overall graph strength did not different between groups. Data are mean ± standard error at 10% edge density. E) Network statistical maps comparing nodal strength and edge weights between CCI day 2 or day 30 and controls (randomization test, 10,000 permutations; blue nodes have p<0.05 and Cohen’s d≥0.08, orange nodes are non-significant, and grey lines p<0.05).

### Analysis of functional network graph properties

Details of network analysis and formal descriptions of graph metrics are published (Pompilus *et al*., 2020). Briefly, weighted matrices were analyzed with Brain Connectivity Toolbox (Rubinov and Sporns, 2010) and MATLAB. Global graph metrics were calculated for edge density thresholds ranging from 2-40%. Node-specific network measures were calculated at a 10% threshold. Node strength was assessed at the individual node level and globally at the graph level to investigate how CCI affected functional connectivity strength. Node strength is the sum of edge weights per node. The average node strength for all edge connected nodes is the network strength.

A probabilistic approach for community detection was applied to calculate a modularity statistic (Q), which indexes the rate of intra-group connections versus connections due to chance (Blondel *et al*., 2008). The procedure starts with a random grouping of nodes and iteratively moving nodes into groups which maximize the value of Q. The final number of modules and node assignments to each group (e.g., community affiliation assignments) was taken as the median of 1000 iterations of the modularity maximization procedure (Pompilus *et al*., 2020). The number of communities per rat, population size of each community, and mean participation coefficient, eigenvector and betweenness centrality of the top four-most populated communities were assessed.

The participation coefficient was used to assess within-module nodes with different levels of participation in connections with nodes in other modules. The calculated participation coefficient (PC) values were classified according to published node cartographic assignments: *PC*<0.05 are ultra-peripheral (UP), 0.05<*PC*<0.62 are peripheral (PC) nodes, 0.62<*PC*<0.80 are non-hub connectors (NHC), and *PC*>0.80 are non-hub kinless (NHK) (Guimera and Amaral, 2005). The presence of highly influential nodes and nodes at the intersection between shortest paths was determined through calculations of betweenness and eigenvector centrality. Nodes with high betweenness centrality are present along many shortest paths between any pair of nodes in a network (Freeman, 1977). Nodes with high eigenvector scores are connected to other high centrality nodes and are considered highly influential within a network (Newman, 2018).

We next analyzed the tendency of assortative vs dissortative mixing of nodes (Newman, 2002). The assortativity index is a Pearson correlation coefficient comparing node strength values between pairs of edge-connected nodes. Positive r values indicate connectivity between pairs of nodes with similar strengths (e.g., high strength nodes pairs with high and low with low), while negative r values indicate cross-pairings between low and high strength nodes. We also generated and analyzed weighted rich club coefficient curves (Φ_w_) at an edge density of 10% (Colizza *et al*., 2006). Nodes belonging to a rich club subnetwork have an above-chance tendency to tightly connect with each other and share the bulk of connections within the network (Colizza *et al*., 2006; van den Heuvel and Sporns, 2011). The approach creates subgraphs containing nodes with strength values at a predetermined degree value, k. For each k-level subgraph, Φ_w_ is calculated as the ratio of the sum of edge weights that connect nodes of degree k or higher (W>k) to the total possible number of summed weights (van den Heuvel and Sporns, 2011):

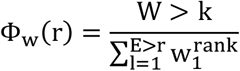

To assess network integration and efficiency, we analyzed the clustering coefficient (CC; an index of the number of connected neighbors of a node) (Onnela *et al*., 2005), characteristic path length (CPL; the lowest average number of edges between node pairs), and the small world coefficient (SWI; which is > 1 for efficient small world networks) (Bassett and Bullmore, 2006). Global efficiency was determined as the average inverse of CPL. Edges were randomly swapped 10 times to generate randomized graphs, preserving original degree and strength distributions (Maslov and Sneppen, 2002; Rubinov and Sporns, 2010). SWI was then calculated as the ratio of lambda to gamma coefficients, where lambda is the ratio of real to random network CC, and gamma the ratio of real to random network CPL (Humphries and Gurney, 2008).

### Functional connectivity network visualizations

Functional connectivity networks were visualized in BrainNet viewer (Xia *et al*., 2013). Center coordinates for each ROI were generated based on parcellations on a previously published rat brain template (Kenkel *et al*., 2016). A 3D whole brain surface mesh file of the rat brain template (in *.byu format) was generated using an image binarization command in FSL (fslmaths) (Jenkinson *et al*., 2002) and mesh construction tools in ITKSNAP (Yushkevich *et al*., 2006). Several 3D brain maps were generated. A first series of connectivity strength maps were produced with the size of nodes (spheres) weighted by node strength and the edges (lines connecting node pairs) weighted by the Pearson coefficient between pairs of nodes (non-thresholded to show all edges at 10% density). Additional maps were generated with node and edge colors representing the module assignment (e.g., community affiliation vector) and node size weighted by eigenvector centrality scores or participation coefficient values.

### Statistical analysis

Statistical analyses and data plotting were carried using MATLAB and GraphPad Prism 9. Two-tailed permutation tests (10,000 randomized inter-group data point swaps) were used to compare data from controls to each of the CCI groups (Zalesky *et al*., 2010; Chung, 2019). Significant differences between groups were considered for p-values < 0.05 and an effect size equal to or greater than 0.8 (large to very large effect size range) (Sawilowsky, 2009). For nodal metrics and pairwise edge comparisons, false discovery rate (FDR) adjustments to p-values were carried out using the Benjamini-Hochberg linear step-up procedure (Benjamini and Hochberg, 1995). We should note, however, that network metrics, particularly across individual spatially distributed nodes, can be highly correlated due to the physical proximity between certain nodes in the brain and because of inherent correlative structure produced by the Pearson r values used as edge weights. Familywise and even FDR p-value corrections in spatial/anatomical data points that rely on correlated measures (e.g., shared signal components across multiple regional data points) can thus significantly reduce power. Thus, FDR adjusted p-values are reported along with uncorrected permutation test p-values and effect sizes, although the latter two criteria for significance are used here. For global graph metrics, p-values for repeated tests over different edge-density thresholds were adjusted using a Bonferroni-Holm (BH) procedure (Holm, 1979; Groppe *et al*., 2011).

## Results

### Structural differences between control and CCI rats

Brain structural differences between control and CCI rats were observed, even after nonlinear warping of each anatomical scan to a reference anatomical template. Figure 2 (top) shows group-averaged normalized (log) Jacobian transformation images of control and CCI day 2 and day 30 rats. Tissue volumetric stretching (expansion) changes relative to the template are noted in yellow-to-red intensity colors and shrinking (contractions) in light-to-dark blue. On average, the control group shows expansion of the cortical mantle along with modest increases in callosal white matter and underlying subcortical (thalamic) areas. The intensity of the observed structural increases differed in CCI rats, particularly at day 2 post-injury. CCI day 2 rats had on average, less cortical expansion, and greater subcortical expansion, with some contraction of white matter structure (splenium of the corpus callosum). Structural differences between CCI day 2 and controls was less intense in CCI day 30 rats (Figure 2, top).

Figure 2 (bottom) shows representative anatomical images of four CCI rats on day 2 and 30, which were registered to the anatomical template. The CCI epicenter sustained significant localized cortical damage across subjects on day 2, which appeared to expand below the cortex to underlying white matter and ventrally towards ipsilateral dorsal hippocampal and thalamic areas, similar to previously reported in histological assessments (Yang *et al*., 2010). The structural differences between the control and CCI groups were partly, but not completely, corrected by the symmetric normalization warping. Areas outside the CCI epicenter, including contralateral structures, appeared well aligned to the template and its accompanying parcellation.

### Contralesional increases in node strength were observed on days 2 and 30 post-CCI

Figure 3B shows 3D functional connectome maps of control (n=23) and CCI groups (day 2 n=23 and day 30 n = 8). Compared to controls, individual nodes on the contralateral hemisphere to the CCI site had greater mean node strength values on both day 2 and 30. Visualization of this effect is enhanced in minimum spanning tree maps in Figure 3C. In addition, reductions in node strength were observed at the ipsilateral CCI area on day 30. When averaged across the entire brain, the local differences in node strength were not significantly different at a network-wide level (Figure 3D). However, differences between control and CCI groups was supported by two-tailed permutation t-tests (Figure 3E). In Figure 3E, blue spheres indicate areas of significant differences in node strength between CCI and control rats (p<0.05, Cohen’s d ≥ 0.8). These areas of significant differences compared to control group were largely present in contra-lateral cortex on day 2 but included ipsilateral structures on day 30 post-CCI. Differences in edges were also investigated between control and CCI groups and these are highlighted as grey connections between nodes in Figure 3E (p<0.05, Cohen’s d ≥ 0.8). We observed a higher density of significantly different edge weights between control and CCI 30 than in day 2 CCI rats.

Node strength values were compared between control and CCI groups at an edge density threshold of 10% (Figure 4). On day 2, CCI rats had greater node strength than controls in the contralateral secondary visual cortex (t=-3.0, p=0.003, FDR=0.08, d=-0.89), ventral subiculum (t=-2.8, p=0.006, FDR=0.07, d=-0.82), globus pallidus (t=-3.0, p=0.003, FDR=0.13, d=-0.89), ventroposterolateral thalamic nucleus (t=-3.0, p=0.004, FDR=0.06, d=-0.87), and lesion nodes 4 and 5 (t=-3.2, p=0.002, FDR=0.16, d=-0.94 and t=-3.0, p=0.003, FDR=0.07, d=-0.89, respectively), which correspond to different subregions of the primary visual cortex (Figure 4A and 4C). On day 2, CCI rats had lower node strength than controls in ipsilateral lateral orbital cortex (t=2.8, p=0.006, FDR=0.08, d=0.82), olfactory tubercle (t=2.7, p=0.007, FDR=0.07, d=0.80), and laterodorsal thalamic nucleus (t=2.8, p=0.005, FDR=0.08, d=0.84) (Figure 4A and 4C).

**Figure 4.**
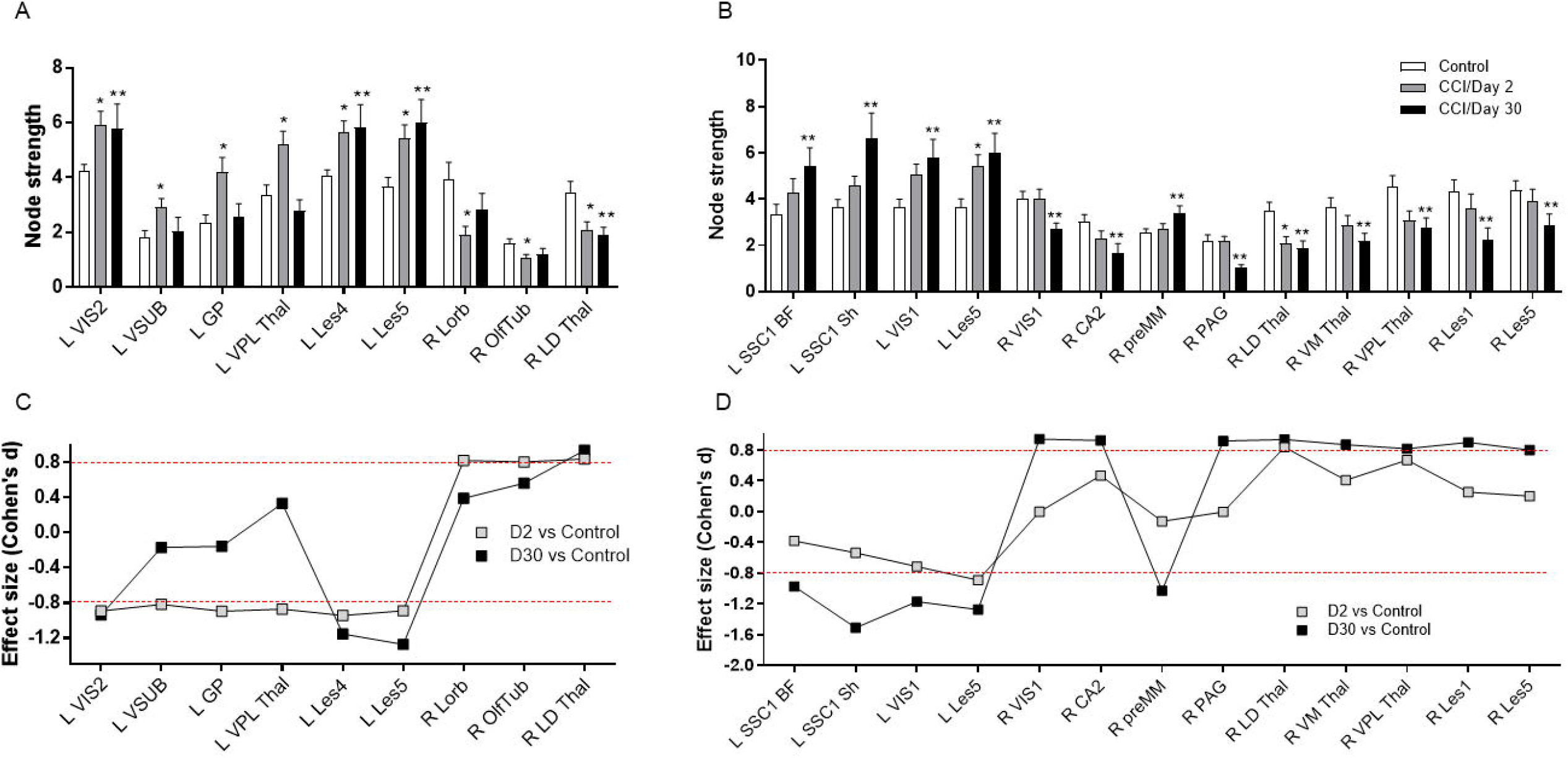
Left hemisphere regions contralateral to the CCI location show increases in node strength at day 2 post-injury, and ipsilateral nodes reductions in node strength. A) Differences are not limited to the cortex and include sensorimotor-projecting thalamic and frontal cortical areas. B) Effect sizes for the differences between control and day 2 CCI are shown in the bottom graph. C) Left hemisphere regions contralateral to the CCI location show increases in node strength at day 30 post-injury, and ipsilateral nodes reductions in node strength. D) Effect sizes for the differences between control and day 30 CCI are shown in the bottom graph. Significant difference between control and CCI day 2* or day 30**, p<0.05 (permutation test) and Cohen’s d ≥ 0.80. Data in bar plots in the top row are mean ± standard error at 10% edge density.

On day 30, CCI rats had greater node strength than controls in contralateral primary somatosensory cortex barrel field (t=-2.3, p=0.04, FDR= 0.30, d=-0.97) and shoulder/limb regions (t=-2.7, p=0.02, FDR= 0.28, d=-1.51), primary visual cortex (t=-2.4, p=0.03, FDR= 0.26, d=-1.17) and lesion node 5, which correspond to a different subregion of primary visual cortex (t=-2.6, p=0.02, FDR= 0.27, d=-1.27). Increased node strength was also observed in ipsilateral premammillary nucleus (t=-2.2, p=0.046, FDR= 0.30, d=-1.03) (Figure 4B and 4D). On day 30, CCI rats had lower node strength than controls in ipsilateral primary visual cortex (t=3.2, p=0.007, FDR= 0.43, d=0.94), hippocampal CA2 (t=2.6, p=0.02, FDR= 0.25, d=0.93), periaqueductal grey (t=3.5, p=0.003, FDR= 0.46, d=0.92), laterodorsal thalamic nucleus (t=3.2, p=0.007, FDR= 0.30, d=0.94), ventromedial thalamic nucleus (t=2.9, p=0.01, FDR= 0.34, d=0.87), ventroposterolateral thalamus (t=2.8, p=0.02, FDR= 0.31, d=0.82), and lesion nodes 1 and 5, which correspond to different subregions of the primary visual cortex (t=2.8, p=0.02, FDR= 0.34, d=0.90; t=2.3, p=0.04, FDR= 0.29, d=0.80) (Figure 4A and 4C). In summary, on day 2 post-injury there are mostly contralateral increases in node strength, whereas both contralateral increases and ipsilateral decreases are observed on day 30.

### Increased modularity and segregation of ‘hub-like’ nodes at 2 days post-injury

In addition to examining the strength of connectivity between individual nodes, we investigated the organization of functional connectivity in control and CCI groups (Figure 5). We observed a significantly greater modularity index in CCI day 2 rats versus controls. This increase was significantly different relative to control group for edge density values of 6%-40% (t=−2.7 to −4.0, p= 0.009 to 0.00001, BH =0.03 to 0.002, d=−0.79 to −1.17). No differences in modularity were observed between controls and CCI rats imaged 30 days post-injury (Figure 5A). In addition, no differences between the groups were observed in the size of the largest module (figure 5B). However, the total number of detected modules was lower in CCI day 2 rats compared to the control group (Figure 5C). This was significant for edge density threshold values of 12% (t=2.7, p=0.009, BH= 0.14, d=0.80), 14% (t=2.8, p=0.005, BH= 0.01, d=0.83), 18% (t=2.8, p=0.007, BH= 0.12, d=0.81), 20% (t=3.6, p=0.0005, BH= 0.01, d=1.06), and 24% (t=2.9, p=0.004, BH= 0.08, d=0.84).

**Figure 5.**
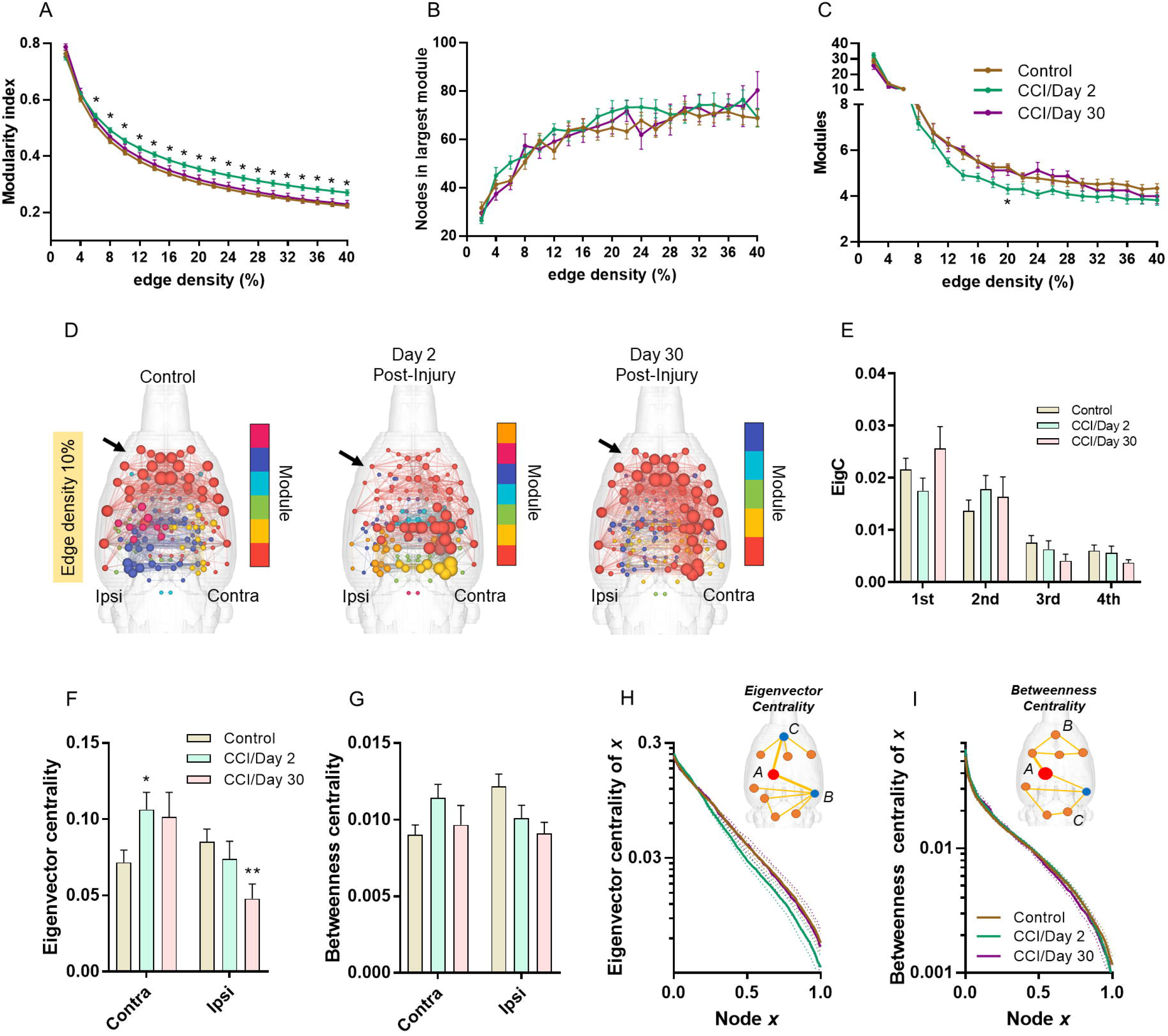
Controlled cortical impact increased network modularity and node eigenvector centrality. A) Modularity index is significantly higher in CCI day 2 post-injury relative to control rats. B) No differences in the size of the largest module. C) The mean number of detected modules was lowest for CCI day 2 post-injury rats. D) 3D connectome maps of node eigenvector centrality across modules. Color coding represents modular group. E) No differences in eigenvector centrality scores across the top four largest modules. F) Compared to control rats, CCI day 2 rats had higher eigenvector centrality and CCI day 30 rats lower scores. H) Eigenvector centrality scores were generally lowest in CCI day 2 rats compared to controls. I) No differences in betweenness centrality were observed. Significant difference between control and CCI day 2* or day 30**, p<0.05 (permutation test) and Cohen’s d ≥ 0.80 (p-values in A and C adjusted using a Bonferroni-Holm procedure). Data in bar plots in F-G are mean ± standard error at 10% edge density.

Figure 5D shows 3D connectome maps with node sizes scaled by eigenvector centrality scores (edge density 10%). In CCI day 2 rats, nodes in the contralateral cortex had the highest eigenvector centrality values (light blue) compared to nodes in the same anatomical location in control rats, whereas bilaterally located nodes in frontal cortical and basal forebrain areas (red nodes) had the lowest. In CCI day 30 rats, nodes in the ipsilateral cortex had the lowest eigenvector centrality values, whereas nodes in frontal cortical and basal forebrain areas were similar to controls. We examined the mean eigenvector centrality values for all nodes contained in each module (Figure 5E). No differences between control and TBI groups were observed in eigenvector centrality scores across the different modules. We next examined the mean eigenvector and betweenness centrality values for all nodes forming part of the ipsi- and contralateral lesion sites (Figure 3A; Figure 5F-I). Contralesional nodes had significantly higher eigenvector scores in CCI day 2 rats compared to controls (t=-2.5, p=0.02, d=-0.72) (Figure 5F). CCI day 30 rats showed a similar but non-significant trend in the contralesional cortex (t=-1.7, p=0.08, d=-0.74). On day 30 post-CCI, rats had lower ipsilateral cortex eigenvector scores compared to controls (t=2.96, p=0.02, d=1.01) (Figure 5F). Eigenvector centrality scores trended lower in CCI day 2 rats compared to control rats although this did not reach statistical significance. No differences in betweenness centrality were observed between the groups (Figure 5G and 5I).

Figure 6A shows 3D connectome maps with node sizes scaled by the participation coefficient value (edge density 10%). In CCI day 2 rats, nodes in frontal cortical and basal forebrain areas had low participation coefficient values compared to the same nodes in control rats (Figure 6A, red colored nodes middle 3D map compared to left 3D map). In CCI day 30 rats, nodal participation coefficient values resembled those of control rats (red colored nodes in 3D map on right). We assessed the mean participation coefficient for all nodes contained in each module (Figure 6B). Compared to controls, nodes in the largest module of CCI day 2 rats had significantly lower participation in cross-modular connectivity (t=3.4, p=0.001, BH=0.006, d=0.99). We also determined the mean participation coefficient for all nodes in the ipsi- and contralateral lesion sites (Figure 6C). Participation coefficient values of contralesional nodes were not different from controls, although a trend towards reduction in CCI day 2 rats was observed (t=1.8, p=0.07, d=0.53) (Figure 6C). Nodes in the ipsilateral cortex, however, had lower participation coefficient values in CCI day 2 rats compared to controls (t=3.1, p=0.005, d=0.90). This difference was not observed on day 30 post-injury. Nodal classification based on the participation coefficient did not differ between groups (Figure 6D).

**Figure 6.**
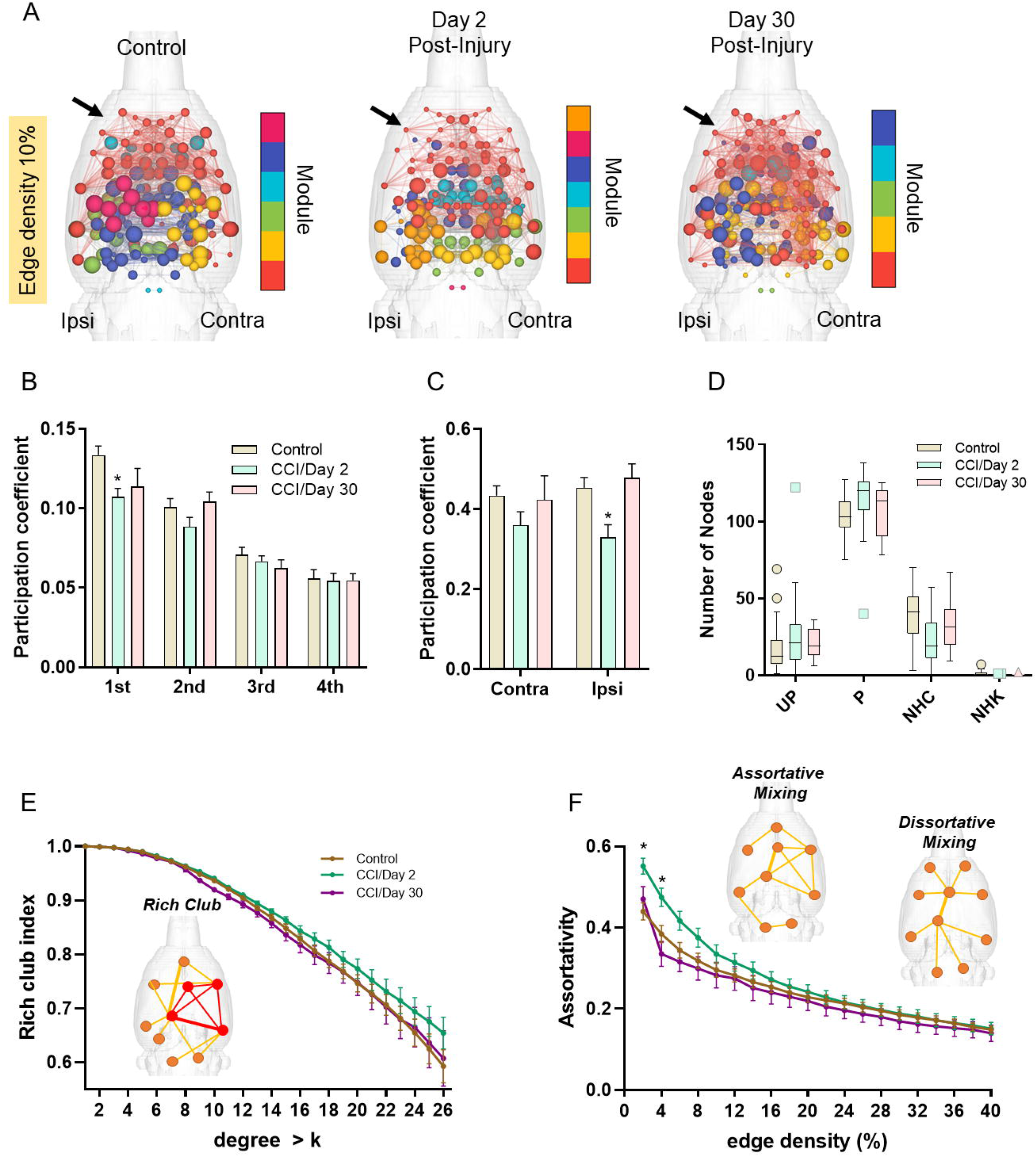
Controlled cortical impact reduced node participation coefficient. A) 3D connectome maps showing node participation coefficient across modules. Color coding represents modular grouping of nodes and node size scaled by participation coefficient. B) Participation coefficient in the largest module (1^st^) is reduced in CCI day 2 post-injury relative to control rats. C) Participation coefficient in the ipsilateral cortex is reduced in CCI day 2 post-injury relative to control rats. D) Control and CCI rats had similar distribution of nodes of distinct cartographic classifications, including ultraperipheral (UP), peripheral (P), non-hub connector (NHC) and non-hub kinless (NHK). E) No differences between rich club index were observed. F) Positive assortativity was higher in CCI day 2 rats compared to controls. Significant difference between control and CCI day 2* or day 30**, p<0.05 (permutation test) and Cohen’s d ≥ 0.80 (p-values in A and C adjusted using a Bonferroni-Holm procedure). Data in bar plots in B-C are mean ± standard error and in data in D is box-whisker plots with 95% confidence bands (at 10% edge density).

Finally, differences in rich club organization and the degree of assortative mixing of nodes were both assessed (Figure 6E and 6F, respectively). Due to the observed increase in contralateral node strength in the TBI groups (Figure 3E) and increased contralateral eigenvector centrality in day 2 CCI rats (Figure 5F), we anticipated similar increases in how high strength nodes interact with other high strength versus low strength nodes. Although we did not observe any group differences in the rich club index (Figure 6E), we observed a significant increase in assortativity index in CCI day 2 rats compared to controls (Figure 6F) (2%: t=-3.9, p=0.0004, BH= 0.007, d=-1.15; 4%: t=-2.9, p=0.005, BH= 0.10, d=-0.86, 6%: t=-2.4, p=0.005, BH= 0.35, d=-0.71). No differences between control and CCI groups in the SWI, global CC and average CPL were observed (Figure 7).

**Figure 7.**
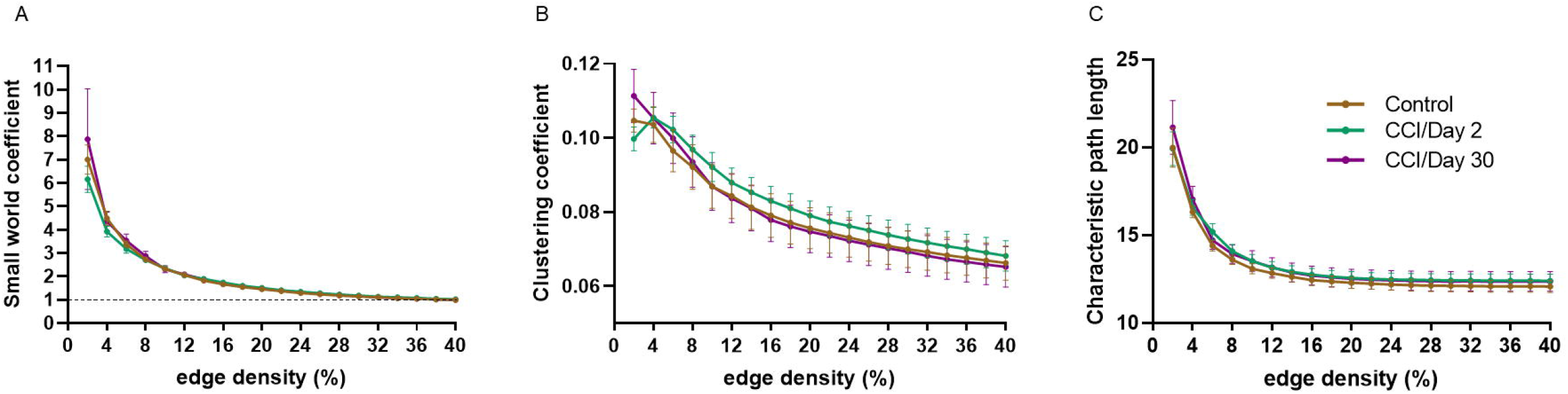
No differences in small world index, clustering coefficient and mean characteristic path length were observed between control and CCI rats. Data are plotted as mean ± standard error at varying edge densities.

## Discussion

Our data indicate that focal injury to the rat cortex resulted in increased node strength in the contralesional hemisphere at days 2 and 30 post-injury. At 30 days post-CCI, we observed significant reductions in node strength in ipsilateral cortex. The brain regions showing node strength differences between CCI and control rats included cortical nodes in or adjacent to the CCI foci (e.g., visual cortical and somatosensory areas) and structures directly underneath and ipsilateral to this region (e.g., thalamic nuclei, hippocampus, PAG) and contralateral homotopic cortical regions.

In addition to node strength, we assessed network modular organization and node centrality. The grouping of nodes into segregated functional units or modules is a salient characteristic of structural and functional brain networks and is highly conserved across various species (Gomez-Gardenes *et al*., 2010; Harriger *et al*., 2012; D’Souza *et al*., 2014; Shih *et al*., 2015; Bardella *et al*., 2016; Zamora-Lopez *et al*., 2016; Brittin *et al*., 2021). This graph metric of network segregation has been shown to be disrupted in TBI patients (Han *et al*., 2014) and is responsive to cognitive training (Han *et al*., 2020). In the present study we observed that rats imaged 2 days after CCI had increased modularity. The CCI day 2 rats had populations of nodes with either low or high eigenvector centrality scores that were segregated into different modules. High centrality scores were observed in modules with nodes located at or near the contralesional cortex. Low centrality scores were observed in modules with nodes anatomically more distant from the injury site. This segregated pattern of low and high centrality nodes is consistent with an observed increase in positive assortative mixing of nodes in CCI day 2 rats. Adding to this segregation of high and low influence nodes was a reduction in node participation in cross-module interactions (reduced participation coefficient) in CCI day 2 rats. These findings suggest that in early stages post-CCI there is transient network topology involving increased functional specialization of separate pools of high and low influence nodes within highly self-contained modules. This network topology appears unfavorable to neuronal communication and to the ‘binding’ of separate streams of information. By day 30 post-injury, nodes in the injury epicenter had lower eigenvector centrality scores compared to control rats, perhaps due to ongoing loss of influence of lesioned structures on overall network topology. However, cross module interactions and assortative mixing did not differ from control rats – suggesting partial recovery.

There are a growing number of clinical TBI studies that use graph theory analytical methods to investigate functional network topology. Nakamura and colleagues were among the first to demonstrate the value of graph theory metrics as potential biomarkers of TBI progression. They imaged 6 subjects over the course of 3- and 6-months post-injury and observed significant reductions in node strength and graph sparsity (Nakamura *et al*., 2009). The small cohort of patients had varying lengths of hospitalization, age, gender, and location of contusions, but all participants had an initial (e.g., 24hr) Glascow Coma Score of 3-8, in the severe range. Compared to controls and to the 6-month timepoint of recovery, at 3-months of recovery TBI patients had shorter CPL, increased global and local efficiency, and reduced small worldness at a graph sparsity of 50% (Nakamura *et al*., 2009). At 6-months post-trauma, mild TBI patients suffering from post-concussion syndrome had reduced modularity compared to controls and this was correlated with symptom severity (Messe *et al*., 2013). Consistent with the present work in rats, patients exposed to blast related mild TBI had lower participation coefficient values compared to non-TBI controls that also experienced blasts (Han *et al*., 2014). Interestingly, reductions in nodal participation in cross-module interactions was most severe within the early timepoint of 90 days post-blast compared to subsequent imaging sessions at 6- and 12-month follow-up (Han *et al*., 2014). This suggests that cross-modular functional connectivity returns over the course of recovery and the participation coefficient calculation is responsive to these network level changes.

At approximately 8 years after injury, chronic TBI patients had significantly lower connectivity within default mode and frontoparietal control networks (DMN and FPCN, respectively), and reduced connectivity of these two networks with the dorsal attention network (DAN) (Han *et al*., 2016). At a graph sparsity threshold greater than in the above cited work by Nakamura and colleagues (Nakamura *et al*., 2009), TBI patients had significantly reduced network efficiency (Han *et al*., 2016). Decreased local efficiency and degree centrality were also reported in mild TBI follow imaging visits at 3 months post-injury, with regions of the DMN being most affected (Sun *et al*., 2021). This is consistent with moderate and severe TBI patients 1-year post-injury, which had lower local integration (lower clustering coefficient) and higher CPL (less efficiency) compared to controls (Gilbert *et al*., 2018), and with mild TBI patients imaged within 10 days which showed increased node strength and CPL (Hou *et al*., 2019). These clinical mild TBI studies agree for the most part that functional connectivity networks of patients have high characteristic path lengths (low efficiency), increased node strength, lower local integration (low clustering). In addition, communication between large functionally specialized groups of nodes (such as DMN, FPCN, and DAN) is disrupted in TBI patients. Our data in the rat CCI model can be considered severe early stage TBI, with a limited analysis across the lifespan of young adult rodents. Nonetheless, our results are in general agreement with increased functional specialization, increased strength of connectivity, and reduced cross module communication in early TBI, which is partially recovered by 30 days. It should be noted that variations across studies may be due to the variability of brain injury across individuals and differences in short- and long-term TBI effects (e.g., location, extent or magnitude of damage, neurological and neuropsychiatric sequelae, age, etc.). In addition, assumptions in the initial steps in network processing such as the choice threshold (graph sparsity) or whether to threshold or not (use individual graph sparsity) can produce different results across laboratories. In some cases, low thresholds that allow the inclusion of weaker edges (much lower Pearson r values) or edges with negative values can lead to spurious results.

Compared to clinical studies, there have been fewer animal fMRI studies of TBI, particularly studies focused on analyzing network interactions following controlled focal or diffuse injury. This might be due to concerns over the inability of fMRI connectivity measurements to disambiguate changes in neurovascular coupling from frank neuronal or synaptic deficits, and valid concerns regarding the confounding effects of anesthetics and sedatives used for functional imaging of rodents. Apart from these important issues, the experimental use of methods that allow well-controlled site-specific lesions and diffuse injuries offers an opportunity to investigate network metrics as potential functional brain biomarkers of TBI. This is illustrated in a within-subjects imaging study in the CCI rat model by Harris and colleagues (Han *et al*., 2016). Rats were imaged in a 7 Tesla MR scanner under continuous medetomidine sedation during baseline conditions (pre-injury), and at 7-, 14- and 28-days post-injury. Their results indicated a significant increase in global node strength, which is consistent with reports of hyperconnectivity in TBI patients (Bernier *et al*., 2017). Consistent with our present findings, CCI rats had significantly increased node strength in contralesional cortex and reduce strength in ipsilateral cortex. In terms of topological changes over time, they report decreases in CPL, increased efficiency, consistent with initial work in TBI patients by Nakamura and colleagues (Nakamura *et al*., 2009). Their data also showed increased global network clustering and transiently reduced modularity index (on day 7 compared to pre-injury), which is consistent with diffusion MRI based tractography studies in the closed head weight drop model in mice (Meningher *et al*., 2020). We did not observe differences in weighted clustering coefficient between the groups (in our between-subjects design), but we did observe a similar but non-significant trend towards increased global transitivity and binarized clustering coefficient in CCI day 2 rats compared to controls (data not shown).

Differences between the Harris et al. study (Han *et al*., 2016) and the present study are perhaps due to the experimental design (e.g., mild to moderate TBI in Harris et al vs a more severe contusion model used in the present study) and/or methods (alpha-2 agonist sedative-hypnotic vs light general anesthesia). Also, the independent use of either a within-subjects vs a between-subjects design limits interpretations between studies. Within-subjects design are highly robust in terms of tracking individual, subject-specific changes over time and over the course of disease progression and recovery. However, experience-dependent changes can also confound interpretations when between-subjects control comparisons are not included in the study. This point is in part illustrated in previous imaging work by Orsini et al (Orsini *et al*., 2018) and Colon-Perez et al (Colon-Perez *et al*., 2019), which used a combination of both within- and between-subjects comparisons and observed changes over the course of several imaging sessions that were separated by weeks. In Orsini et al (Orsini *et al*., 2018), while drug treatment was observed to lead to increases in functional connectivity (when comparing pre- to post), these changes were not specific to the drug and were observed with a natural reward given for the same amount of time. In addition, baseline control groups receiving no treatment over the course of the 3 imaging sessions still showed differences when compared to baseline imaging session (Orsini *et al*., 2018). In Colon-Perez et al (Colon-Perez *et al*., 2019), the inclusion of an untrained group of rats functionally imaged at the same timepoints as aged and young rats that underwent cognitive training was necessary to demonstrate that in fact training increased rich club organization and node strength (Colon-Perez *et al*., 2019). Thus, there are gains in statistical power by including a mixed between/within-subjects multisession imaging design in TBI research in animal models. There have been other significant functional imaging studies in animal models of TBI. These have focused on traditional seed-based functional connectivity but did not examine functional network topology (Heffernan *et al*., 2013; Mishra *et al*., 2014; Kulkarni *et al*., 2019).

Cellular and neurophysiological links to stimulus-based BOLD hyperexcitability in contralesional cortex of CCI rats have been previously reported (Verley *et al*., 2018). Conversely, Hypoexcitability in ipsilateral cortex has also been demonstrated for up to 56 days in the FPI rat model (Niskanen *et al*., 2013). Glutamatergic hyperexcitability involved in TBI-induced epileptiform activity was shown in the CCI injured rat cortex (at 2-4 weeks post-injury) and was observed to spread to adjacent regions, an effect associated with loss of GABAergic control (Yang *et al*., 2010; Cantu *et al*., 2015). In mice, CCI reduced GABAergic cell counts in the ipsilateral dorsal and medial hilar of the hippocampal dentate gyrus and reduced inhibitory post-synaptic currents were observed as early as 1 week and for as long as 13 weeks (Butler *et al*., 2016). Under control conditions (non-TBI), transcallosal fibers, which in adults are almost exclusively comprised of glutamatergic projections, exert inhibitory control over contralateral primary sensory receptor fields (Clarey *et al*., 1996; Pascual-Leone *et al*., 2005; Pluto *et al*., 2005; Schafer *et al*., 2012). Ipsilateral reductions in neuronal activity have been observed to widen contralateral neocortical receptive fields, possibly by reducing transcallosal cross-hemispheric inhibitory control (Clarey *et al*., 1996). Contralateral cortical BOLD increases in response to limb stimulation in rats is accompanied by ipsilateral BOLD signal reductions (Devor *et al*., 2008). These separate lines of evidence suggests that reductions in the influence of neuronal populations in the ipsi-lesioned cortex can remove these transcallosal inhibitory control inputs over the contralateral cortex. The microcircuit mechanism may involve transcallosal glutamatergic projections that synapse onto contralaterally located GABAergic neurons. While most callosal inputs terminate in layer 2/3 pyramidal neurons, these also terminate on soma and dendrites of parvalbumin positive GABA interneurons in deep neocortical layer 6 (Karayannis *et al*., 2007). These neurons are thought to play a key role in synchronization of thalamocortical loops (Karayannis *et al*., 2007). Loss of transcallosal glutamatergic inputs may therefore lead to decrease excitability of GABA cells and result in pyramidal overexcitation of the contralateral cortex. This could partly explain the observed compensatory increases in functional connectivity and reorganization of hub nodes. Spontaneous outputs from these disinhibited pyramidal cells can lead to downstream plasticity related changes in neuronal activity, as observed in peripheral sensory deprivation models (Pluto *et al*., 2005; Petrus *et al*., 2020).

We should indicate that although neocortical changes in GABA and glutamatergic excitability can play a role in brain wide changes in functional connectivity in TBI, mechanisms involved in neurovascular coupling are also likely important (Ahmad *et al*., 2012; Sakai *et al*., 2021; Wu *et al*., 2021). Loss of synchrony in BOLD could reflect underlying changes in neurovascular coupling, particularly with the occurrence of gliosis and astrocytic changes in the injury epicenter. The neuronal and cerebrovascular/hemodynamic mechanisms contributing to the observed increases in contralesional node strength, and the alterations in modularity and ‘hub’ node organization, warrant future investigations. Network assessments in TBI animal models opens the door to establishing mechanistic links between graph theory metrics and underlying neuronal, glial, and vascular mechanisms, and behavior. Understanding the mechanisms of these and other reported connectomic changes following TBI can help with therapeutic interventions that use functional connectomic readouts as indices (biomarkers) of recovery or treatment efficacy.

## Acknowledgments

This study was funded by a National Institute on Neurological Disorders and Stroke grant (UG3 NS106938) to KKW. The contents of this manuscript are solely the responsibility of the authors and do not necessarily represent the official views of the funding agencies. This work was performed in the McKnight Brain Institute at the National High Magnetic Field Laboratory’s AMRIS Facility, which is supported by National Science Foundation Cooperative Agreement No. DMR-1644779 and the State of Florida.

## References

Ahmad A, Crupi R, Impellizzeri D, Campolo M, Marino A, Esposito E, et al. Administration of palmitoylethanolamide (PEA) protects the neurovascular unit and reduces secondary injury after traumatic brain injury in mice. Brain Behav Immun 2012; 26(8): 1310–21.

Arciniegas DB. Addressing neuropsychiatric disturbances during rehabilitation after traumatic brain injury: current and future methods. Dialogues Clin Neurosci 2011; 13(3): 325–45.

Bardella G, Bifone A, Gabrielli A, Gozzi A, Squartini T. Hierarchical organization of functional connectivity in the mouse brain: a complex network approach. Sci Rep 2016; 6: 32060.

Bassett DS, Bullmore E. Small-world brain networks. Neuroscientist 2006; 12(6): 512–23.

Benjamini Y, Hochberg Y. Controlling the false discovery rate: a practical and powerful approach to multiple testing. Journal of the Royal Statistical Society B 1995; 57: 289–300.

Bernier RA, Roy A, Venkatesan UM, Grossner EC, Brenner EK, Hillary FG. Dedifferentiation Does Not Account for Hyperconnectivity after Traumatic Brain Injury. Front Neurol 2017; 8: 297.

Blondel VD, Guillaume JL, Lambiotte R, Lefebvre E. Fast unfolding of communities in large networks. Journal of Statistical Mechanics: Theory and Experiment 2008: 1–12.

Brittin CA, Cook SJ, Hall DH, Emmons SW, Cohen N. A multi-scale brain map derived from whole-brain volumetric reconstructions. Nature 2021; 591(7848): 105–10.

Butler CR, Boychuk JA, Smith BN. Differential effects of rapamycin treatment on tonic and phasic GABAergic inhibition in dentate granule cells after focal brain injury in mice. Exp Neurol 2016; 280: 30–40.

Cantu D, Walker K, Andresen L, Taylor-Weiner A, Hampton D, Tesco G, et al. Traumatic Brain Injury Increases Cortical Glutamate Network Activity by Compromising GABAergic Control. Cereb Cortex 2015; 25(8): 2306–20.

Carron SF, Yan EB, Alwis DS, Rajan R. Differential susceptibility of cortical and subcortical inhibitory neurons and astrocytes in the long term following diffuse traumatic brain injury. J Comp Neurol 2016; 524(17): 3530–60.

CDC. Surveillance of TBI-related emergency department visits, hospitalizations, and deaths - United States, 2001-2010. U.S.: National Center for Injury Prevention and Control; 2019.

Chou N, Wu J, Bai Bingren J, Qiu A, Chuang KH. Robust automatic rodent brain extraction using 3-D pulse-coupled neural networks (PCNN). IEEE Trans Image Process 2011; 20(9): 2554–64.

Chung MK. Brain Network Analysis: Cambridge University Press; 2019.

Clarey JC, Tweedale R, Calford MB. Interhemispheric modulation of somatosensory receptive fields: evidence for plasticity in primary somatosensory cortex. Cereb Cortex 1996; 6(2): 196206.

Colizza V, Flammini A, Serrano MA, Vespignani A. Detecting rich-club ordering in complex networks. Nature Physics 2006; 2: 110–5.

Colon-Perez LM, Turner SM, Lubke KN, Pompilus M, Febo M, Burke SN. Multiscale Imaging Reveals Aberrant Functional Connectome Organization and Elevated Dorsal Striatal Arc Expression in Advanced Age. eNeuro 2019; 6(6).

Cox RW. AFNI: software for analysis and visualization of functional magnetic resonance neuroimages. Comput Biomed Res 1996; 29(3): 162–73.

D’Souza DV, Jonckers E, Bruns A, Kunnecke B, von Kienlin M, Van der Linden A, et al. Preserved modular network organization in the sedated rat brain. PLoS One 2014; 9(9): e106156.

De Simoni S, Jenkins PO, Bourke NJ, Fleminger JJ, Hellyer PJ, Jolly AE, et al. Altered caudate connectivity is associated with executive dysfunction after traumatic brain injury. Brain 2018; 141(1): 148–64.

Devor A, Hillman EM, Tian P, Waeber C, Teng IC, Ruvinskaya L, et al. Stimulus-induced changes in blood flow and 2-deoxyglucose uptake dissociate in ipsilateral somatosensory cortex. J Neurosci 2008; 28(53): 14347–57.

Fagerholm ED, Hellyer PJ, Scott G, Leech R, Sharp DJ. Disconnection of network hubs and cognitive impairment after traumatic brain injury. Brain 2015; 138(Pt 6): 1696–709.

Freeman LC. A set of measures of centrality based on betweenness. Sociometry 1977; 40: 3541.

Gilbert N, Bernier RA, Calhoun VD, Brenner E, Grossner E, Rajtmajer SM, et al. Diminished neural network dynamics after moderate and severe traumatic brain injury. PLoS One 2018; 13(6): e0197419.

Gomez-Gardenes J, Zamora-Lopez G, Moreno Y, Arenas A. From modular to centralized organization of synchronization in functional areas of the cat cerebral cortex. PLoS One 2010; 5(8): e12313.

Groppe DM, Urbach TP, Kutas M. Mass univariate analysis of event-related brain potentials/fields I: a critical tutorial review. Psychophysiology 2011; 48(12): 1711–25.

Guimera R, Amaral LAN. Cartography of complex networks: Modules and universal roles. Journal of Statistical Mechanics: Theory and Experiment 2005: 1–13.

Hall ED, Bryant YD, Cho W, Sullivan PG. Evolution of post-traumatic neurodegeneration after controlled cortical impact traumatic brain injury in mice and rats as assessed by the de Olmos silver and fluorojade staining methods. J Neurotrauma 2008; 25(3): 235–47.

Hall ED, Sullivan PG, Gibson TR, Pavel KM, Thompson BM, Scheff SW. Spatial and temporal characteristics of neurodegeneration after controlled cortical impact in mice: more than a focal brain injury. J Neurotrauma 2005; 22(2): 252–65.

Han K, Chapman SB, Krawczyk DC. Disrupted Intrinsic Connectivity among Default, Dorsal Attention, and Frontoparietal Control Networks in Individuals with Chronic Traumatic Brain Injury. J Int Neuropsychol Soc 2016; 22(2): 263–79.

Han K, Chapman SB, Krawczyk DC. Cognitive Training Reorganizes Network Modularity in Traumatic Brain Injury. Neurorehabil Neural Repair 2020; 34(1): 26–38.

Han K, Mac Donald CL, Johnson AM, Barnes Y, Wierzechowski L, Zonies D, et al. Disrupted modular organization of resting-state cortical functional connectivity in U.S. military personnel following concussive ‘mild’ blast-related traumatic brain injury. Neuroimage 2014; 84: 76–96.

Harriger L, van den Heuvel MP, Sporns O. Rich club organization of macaque cerebral cortex and its role in network communication. PLoS One 2012; 7(9): e46497.

Harris NG, Verley DR, Gutman BA, Thompson PM, Yeh HJ, Brown JA. Disconnection and hyper-connectivity underlie reorganization after TBI: A rodent functional connectomic analysis. Exp Neurol 2016; 277: 124–38.

Heffernan ME, Huang W, Sicard KM, Bratane BT, Sikoglu EM, Zhang N, et al. Multi-modal approach for investigating brain and behavior changes in an animal model of traumatic brain injury. J Neurotrauma 2013; 30(11): 1007–12.

Hegde S. Music-based cognitive remediation therapy for patients with traumatic brain injury. Front Neurol 2014; 5: 34.

Hillary FG, Slocomb J, Hills EC, Fitzpatrick NM, Medaglia JD, Wang J, et al. Changes in resting connectivity during recovery from severe traumatic brain injury. Int J Psychophysiol 2011; 82(1): 115–23.

Holm S. A simple rejective multiple test procedure. Scandinavian Journal of Statistics 1979; 6(2): 65–70.

Hou W, Sours Rhodes C, Jiang L, Roys S, Zhuo J, JaJa J, et al. Dynamic Functional Network Analysis in Mild Traumatic Brain Injury. Brain Connect 2019; 9(6): 475–87.

Humphries MD, Gurney K. Network ‘small-world-ness’: a quantitative method for determining canonical network equivalence. PLoS One 2008; 3(4): e0002051.

Jenkinson M, Bannister P, Brady M, Smith S. Improved optimization for the robust and accurate linear registration and motion correction of brain images. Neuroimage 2002; 17(2): 825–41.

Johnstone VP, Wright DK, Wong K, O’Brien TJ, Rajan R, Shultz SR. Experimental Traumatic Brain Injury Results in Long-Term Recovery of Functional Responsiveness in Sensory Cortex but Persisting Structural Changes and Sensorimotor, Cognitive, and Emotional Deficits. J Neurotrauma 2015; 32(17): 1333–46.

Karayannis T, Huerta-Ocampo I, Capogna M. GABAergic and pyramidal neurons of deep cortical layers directly receive and differently integrate callosal input. Cereb Cortex 2007; 17(5): 1213–26.

Kenkel WM, Yee JR, Moore K, Madularu D, Kulkarni P, Gamber K, et al. Functional magnetic resonance imaging in awake transgenic fragile X rats: evidence of dysregulation in reward processing in the mesolimbic/habenular neural circuit. Transl Psychiatry 2016; 6: e763.

Klein A, Andersson J, Ardekani BA, Ashburner J, Avants B, Chiang MC, et al. Evaluation of 14 nonlinear deformation algorithms applied to human brain MRI registration. Neuroimage 2009; 46(3): 786–802.

Kulkarni P, Morrison TR, Cai X, Iriah S, Simon N, Sabrick J, et al. Neuroradiological Changes Following Single or Repetitive Mild TBI. Front Syst Neurosci 2019; 13: 34.

Maslov S, Sneppen K. Specificity and stability in topology of protein networks. Science 2002; 296(5569): 910–3.

Meningher I, Bernstein-Eliav M, Rubovitch V, Pick CG, Tavor I. Alterations in Network Connectivity after Traumatic Brain Injury in Mice. J Neurotrauma 2020; 37(20): 2169–79.

Messe A, Caplain S, Pelegrini-Issac M, Blancho S, Levy R, Aghakhani N, et al. Specific and evolving resting-state network alterations in post-concussion syndrome following mild traumatic brain injury. PLoS One 2013; 8(6): e65470.

Mishra AM, Bai X, Sanganahalli BG, Waxman SG, Shatillo O, Grohn O, et al. Decreased resting functional connectivity after traumatic brain injury in the rat. PLoS One 2014; 9(4): e95280.

Nakamura T, Hillary FG, Biswal BB. Resting network plasticity following brain injury. PLoS One 2009; 4(12): e8220.

Newman ME. Assortative mixing in networks. Phys Rev Lett 2002; 89(20): 208701.

Newman ME. Networks: an introduction. 2nd Edition ed. New York: Oxford University Press; 2018.

Niskanen JP, Airaksinen AM, Sierra A, Huttunen JK, Nissinen J, Karjalainen PA, et al. Monitoring functional impairment and recovery after traumatic brain injury in rats by FMRI. J Neurotrauma 2013; 30(7): 546–56.

Onnela JP, Saramaki J, Kertesz J, Kaski K. Intensity and coherence of motifs in weighted complex networks. Phys Rev E Stat Nonlin Soft Matter Phys 2005; 71(6 Pt 2): 065103.

Orsini CA, Colon-Perez LM, Heshmati SC, Setlow B, Febo M. Functional Connectivity of Chronic Cocaine Use Reveals Progressive Neuroadaptations in Neocortical, Striatal, and Limbic Networks. eNeuro 2018; 5(4).

Pascual-Leone A, Amedi A, Fregni F, Merabet LB. The plastic human brain cortex. Annu Rev Neurosci 2005; 28: 377–401.

Petrus E, Dembling S, Usdin T, Isaac JTR, Koretsky AP. Circuit-Specific Plasticity of Callosal Inputs Underlies Cortical Takeover. J Neurosci 2020; 40(40): 7714–23.

Ping X, Jin X. Transition from Initial Hypoactivity to Hyperactivity in Cortical Layer V Pyramidal Neurons after Traumatic Brain Injury In Vivo. J Neurotrauma 2016; 33(4): 354–61.

Pluto CP, Chiaia NL, Rhoades RW, Lane RD. Reducing contralateral SI activity reveals hindlimb receptive fields in the SI forelimb-stump representation of neonatally amputated rats. J Neurophysiol 2005; 94(3): 1727–32.

Pompilus M, Colon-Perez LM, Grudny MM, Febo M. Contextual experience modifies functional connectome indices of topological strength and efficiency. Sci Rep 2020; 10(1): 19843.

Rubinov M, Sporns O. Complex network measures of brain connectivity: uses and interpretations. Neuroimage 2010; 52(3): 1059–69.

Sakai K, Takata F, Yamanaka G, Yasunaga M, Hashiguchi K, Tominaga K, et al. Reactive pericytes in early phase are involved in glial activation and late-onset hypersusceptibility to pilocarpine-induced seizures in traumatic brain injury model mice. J Pharmacol Sci 2021; 145(1): 155–65.

Sawilowsky SS. New effect size rules of thumb. Journal of Modern Applied Statistical Methods 2009; 8(2): 597–9.

Schafer K, Blankenburg F, Kupers R, Gruner JM, Law I, Lauritzen M, et al. Negative BOLD signal changes in ipsilateral primary somatosensory cortex are associated with perfusion decreases and behavioral evidence for functional inhibition. Neuroimage 2012; 59(4): 3119–27.

Sharp DJ, Beckmann CF, Greenwood R, Kinnunen KM, Bonnelle V, De Boissezon X, et al. Default mode network functional and structural connectivity after traumatic brain injury. Brain 2011; 134(Pt 8): 2233–47.

Shih CT, Sporns O, Yuan SL, Su TS, Lin YJ, Chuang CC, et al. Connectomics-based analysis of information flow in the Drosophila brain. Curr Biol 2015; 25(10): 1249–58.

Sun Y, Wang S, Gan S, Niu X, Yin B, Bai G, et al. Serum neuron-specific enolase levels associated with connectivity alterations in anterior default mode network after mild traumatic brain injury. J Neurotrauma 2021.

Thomas TC, Colburn TA, Korp K, Khodadad A, Lifshitz J. Translational Considerations for Behavioral Impairment and Rehabilitation Strategies after Diffuse Traumatic Brain Injury. In: Kobeissy FH, editor. Brain Neurotrauma: Molecular, Neuropsychological, and Rehabilitation Aspects. Boca Raton (FL); 2015.

Tustison NJ, Avants BB, Cook PA, Zheng Y, Egan A, Yushkevich PA, et al. N4ITK: improved N3 bias correction. IEEE Trans Med Imaging 2010; 29(6): 1310–20.

van den Heuvel MP, Sporns O. Rich-club organization of the human connectome. J Neurosci 2011; 31(44): 15775–86.

Verley DR, Torolira D, Pulido B, Gutman B, Bragin A, Mayer A, et al. Remote Changes in Cortical Excitability after Experimental Traumatic Brain Injury and Functional Reorganization. J Neurotrauma 2018; 35(20): 2448–61.

Wu J, Li H, He J, Tian X, Luo S, Li J, et al. (Downregulation of microRNA-9-5p promotes synaptic remodeling in the chronic phase after traumatic brain injury). Cell Death Dis 2021; 12(1): 9.

Xia M, Wang J, He Y. BrainNet Viewer: a network visualization tool for human brain connectomics. PLoS One 2013; 8(7): e68910.

Yang L, Afroz S, Michelson HB, Goodman JH, Valsamis HA, Ling DS. Spontaneous epileptiform activity in rat neocortex after controlled cortical impact injury. J Neurotrauma 2010; 27(8): 15418.

Yang Z, Wang P, Morgan D, Lin D, Pan J, Lin F, et al. Temporal MRI characterization, neurobiochemical and neurobehavioral changes in a mouse repetitive concussive head injury model. Sci Rep 2015; 5: 11178.

Yushkevich PA, Piven J, Hazlett HC, Smith RG, Ho S, Gee JC, et al. User-guided 3D active contour segmentation of anatomical structures: significantly improved efficiency and reliability. Neuroimage 2006; 31(3): 1116–28.

Zalesky A, Fornito A, Harding IH, Cocchi L, Yucel M, Pantelis C, et al. Whole-brain anatomical networks: does the choice of nodes matter? Neuroimage 2010; 50(3): 970–83.

Zamora-Lopez G, Chen Y, Deco G, Kringelbach ML, Zhou C. Functional complexity emerging from anatomical constraints in the brain: the significance of network modularity and rich-clubs. Sci Rep 2016; 6: 38424.

